# Functional Annotation of Novel Heat Stress-responsive Genes in Rice Utilizing Public Transcriptomes and Structurome

**DOI:** 10.1101/2025.09.29.679423

**Authors:** Sora Yonezawa, Hidemasa Bono

## Abstract

**Motivation:** Life science databases include large collections of public transcriptome and large-scale structural data. The reuse and integration of these datasets may facilitate the identification of understudied genes and enable functional annotation across distantly related species, including plants and humans.

**Results:** In this study, we used heat stress-responsive genes in rice as a model to functionally annotate previously understudied genes by integrating publicly available transcriptome data with structural information from the AlphaFold Protein Structure Database. Initially, we conducted a meta-analysis of public heat stress-related transcriptome datasets, identified gene groups, and verified stress-related terms through enrichment analysis. Subsequently, we performed structural alignment and sequence alignment between rice and human proteins, focusing on candidates exhibiting low sequence similarity but high structural similarity (LS–HS conditions). We further incorporated supplemental data from public databases, including shared domain information between rice and human. This approach yielded a unique set of LS–HS candidates, notably those associated with metal homeostasis, such as iron and copper metabolism. Overall, our integrative method provided insights into these genes by leveraging diverse, publicly available datasets.

**Availability and implementation:** The “plant2human workflow” for this analysis is available at https://doi.org/10.48546/WORKFLOWHUB.WORKFLOW.1206.8.

## 1 Introduction

Public databases in life sciences host diverse types of data and vast amounts of information. The National Genomics Data Center Database Commons, a comprehensive repository that curates life science databases, comprises over 7,300 life science-related databases across 13 categories, including “Gene genome and annotation,” “Health and medicine,” and “Expression” (accessed September 12, 2025) [1]. Access to these data resources supports the development of biological research and analyses.

Among these resources, transcriptome data have expanded rapidly due to the widespread adoption of next-generation sequencing. For example, the Gene Expression Omnibus (GEO) at the US National Center for Biotechnology Information (NCBI) is a public database that stores gene expression and epigenomic data; currently, the “Expression profiling by high throughput sequencing” category contains over 120,000 series (https://www.ncbi.nlm.nih.gov/geo/summary/?type=series) (accessed September 13, 2025) [2]. NCBI GEO data can be reused for various purposes, including the integration of different datasets to identify new biological trends [2]. We previously conducted a meta-analysis of human and mouse heat-stressed transcriptome data from NCBI GEO, integrating transcription factor binding and literature data. This analysis identified heat stress-related genes that are relatively understudied compared to heat shock proteins (HSPs) [3]. This approach can be applied to plant research. For example, meta-analyses of publicly available RNA-sequencing (RNA-Seq) datasets have been conducted in *Arabidopsis thaliana* to identify stress-responsive genes under various stress conditions [4]. Hence, the reuse of public transcriptomic data, particularly through meta-analysis, facilitates new approaches to gene exploration.

As the volume of data accumulated in public databases has increased, new types of massive data have emerged. The AlphaFold Protein Structure Database (AFDB) has become a notable resource, containing more than 214 million predicted protein structures, including those of plants [5], [6]. As a result, it has become feasible to investigate biological questions through the lens of the large-scale protein “structurome.” Studies using the AFDB have explored novel protein structure clusters and domain information, yielding novel biological insights [7], [8].

Quantitative analyses have shown that structural regions within the core regions of conserved protein domains are three to ten times more conserved than their sequences [9], which broadens the range of genes available for comparative and functional annotation, including between distantly related species, such as plants and humans. Functional annotation of plant and human genes has been documented in several studies. In *Arabidopsis*, research has confirmed the function of genes similar to the human *BRCA2* (BRCA2 DNA repair associated) gene [10]. Furthermore, investigations into genes orthologous to human disease-related genes in *Arabidopsis* and *Populus trichocarpa* suggest the potential applicability of plant models in human disease research [11]. However, approaches primarily based on sequence similarity may overlook genes with limited sequence conservation. Overall, the conservation of protein structure and the availability of large-scale structural data could expand the scope for identifying relationships among distantly related species, supporting further cross-species comparisons and annotations.

The expansion of public transcriptome data and the emergence of the structurome (AFDB) reflect the broadening scope of publicly available resources in the life sciences. Recently, the concept of “Unknome” has been proposed. The research group that proposed the Unknome identified clusters using the “knownness score,” a metric derived from Gene Ontology (GO) annotation evidence codes from the Universal Protein Resource (UniProt), and subsequently established the Unknome database. Genes within clusters with low knownness scores represent those that have received limited attention. Genes with low knownness scores that were conserved in humans and *Drosophila melanogaster* and present in at least 80% of the available metazoan genomes were selected and functionally validated via RNAi in *D. melanogaster*. These genes may be involved in essential functions, including stress resistance [12]. Therefore, leveraging public databases to identify understudied genes can facilitate their discovery and support further biological research.

In this study, public transcriptomes and structural data were integrated to develop an analytical workflow for identifying understudied genes and facilitating functional annotation. As a proof of concept, public rice (*Oryza sativa*) transcriptomes related to heat stress were mapped to the AFDB structure database. Rice serves as a suitable model organism for this purpose because of its comprehensive public transcriptome datasets, especially concerning heat stress. Additionally, HSPs, which play an important role of the rice heat stress response [13], [14], are widely conserved at the protein sequence level across distantly related species, including humans; HSP90 is one of the representative example [15].Therefore, this approach was used to identify heat stress-responsive genes that, while lacking sequence similarity, share similar protein structures. By comparing these findings to human structural data, we identified understudied genes and conducted functional annotations based on existing human knowledge.

## 2 Methods

### 2.1. Curation of Heat Stress-related Rice Transcriptome Data

We curated heat stress-related rice (*O. sativa*) RNA-Seq datasets from NCBI GEO [2] using the following search query: ((((((((((heat stress) OR heat shock) OR heat shock response) OR high temperature) OR high temperature stress) OR heat-sensitive) OR heat stress tolerance) OR high night temperature) AND “Oryza sativa”[porgn: txid 4530])) AND “expression profiling by high throughput sequencing”[Filter]) (accessed January 20, 2024). Data, including japonica and indica subspecies, were manually curated based on the described experimental conditions and complete data for heat stress and control conditions.

### 2.2. Transcriptome Data Processing and Quantification

Fasterq-dump from the NCBI SRA Toolkit (ver. 3.0.1) [16], fastp (ver. 0.23.4) [17], and salmon (ver. 1.10.2) [18] was used for FASTQ file retrieval, trimming, and expression quantification. For expression quantification in salmon, the japonica subspecies transcript FASTA file was retrieved from Ensembl Plants release 58 (accessed February 1, 2024) [19] as a reference transcript. Gene expression levels were summarized using tximport software (Bioconductor package release ver. 3.19) [20] and calculated as scaled transcripts per million (scaled TPM).

### 2.3. Calculation of Expression Ratio and HN-score

Following our previously described study protocol [3], we defined the HN-score (Heat-stress and Non-treatment scores) to capture the gene-level responses to heat stress. The calculation procedure is summarized below:

#### 2.3.1 Calculation of expression ratio (HN-ratio)

The heat stress and non-treatment condition data were paired, and expression ratios (HN-ratio) were calculated for all genes using the following Equation (1):

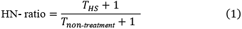

where *T*_*HS*_ and *T*_*non-treatment*_ are the scaled TPM for each gene under heat stress and non-treatment condition pairs, respectively.

#### 2.3.2 Gene classification based on HN-ratio and calculation of HN-score

Based on the HN-ratio and defined 5-fold and 1/5-fold thresholds, all genes were classified as “upregulated,” “downregulated,” or “unchanged.” For instance, if the HN-ratio of Gene A in a given heat stress and non-treatment pair was 5, it was considered “upregulated,” while if the HN-ratio for Gene B was 1/5, it was considered “downregulated.” If Gene C did not meet either threshold, it was considered “unchanged.”

The counts in the three categories were aggregated across all the pairs. The HN-score was calculated as the difference between the “upregulated” and “downregulated” counts for each gene. The top and bottom genes were selected based on their HN-scores and used for subsequent analyses. Detailed annotation information for these selected genes was obtained from the Rice Annotation Project Database (version IRGSP-1.0 2025-03-19) (accessed July 25, 2025) [21].

### 2.4. Enrichment Analysis and Semantic Similarity Analysis

Enrichment analysis was performed using GOAtools (ver. 1.4.12) [22] with GO annotation data obtained from Ensembl Plants release 58 on a selected group of rice genes. Obsolete GO terms were replaced with terms from the latest GO resources (release: March 6, 2025) [23]. The semantic similarity among the enriched terms was calculated using simona (ver.1.4.0) [24] with the “Sim_WP_1994” method [25]. The similarities of all term pairs were visualized as clusters using the SimplifyEnrichment package (ver. 2.0.0) [26].

### 2.5. Structural Similarity and Sequence Similarity Comparison

#### 2.5.1. Structural prediction data retrieval

UniProt accessions corresponding to the IDs of the rice genes selected based on the HN-score were retrieved using Unipressed (ver. 1.3.0) [27] and UniProt Web Application Programming Interface (API) (accessed May 23, 2025) [28]. Macromolecular crystallographic information files (mmCIFs) of the predicted protein structures corresponding to the UniProt accessions were retrieved using the AFDB (version 4) Web API (accessed May 23, 2025) [5].

#### 2.5.2. Structural similarity search

Foldseek (ver. 9-427df8a) was used for the structural analyses [29]. The AFDB database, which contains approximately 214 million entries, was indexed using the foldseek databases command [6]. The collected rice mmCIF files were queried against the AFDB using the foldseek easy-search command. Two alignment modes were applied for structural alignment: (1) 3Di+AA Goto–Smith–Waterman local alignment (3Di+AA) and (2) TM-alignment (Foldseek-TM), which emphasizes global structural features [29]. From the resulting cross-species hits, rice–human hit pairs were extracted for downstream analysis.

#### 2.5.3. Sequence similarity

To obtain protein sequence similarity for the same set of hit pairs, the NCBI Basic Local Alignment Search Tool (BLAST) (ver.2.16.0) [30] and European Molecular Biology Open Software Suite (EMBOSS) package (ver. 6.5.7) were used [31]. All protein sequences used for structure prediction were retrieved from the AFDB FTP site (accessed August 9, 2025) [6] and indexed using the makeblastdb command. Rice and human protein sequences identified via structural alignment were retrieved in the multi-FASTA format using the blastdbcmd command. The multi-FASTA files were split with EMBOSS seqretsplit, followed by pairwise alignments of rice and human protein sequences using EMBOSS Needle for global alignment and EMBOSS Water for local alignment.

#### 2.5.4. Filtering criteria

To focus on global structural similarity, minimize redundant matches, and ensure a clear gene-level analysis of human data, the following filters were applied to rice–human Foldseek hit pairs:

##### (1) Alignment coverage

Coverage of 50% or more on both proteins to prioritize global over local alignments.

##### (2) Redundancy removal within each rice gene

When multiple rice UniProt accessions from the same gene were aligned to the same human structure, the alignment with the highest coverage was retained, and ties were broken using the average Local Distance Difference Test (lDDT) score [32].

##### (3) Human cross-reference

Only hit pairs with human UniProt accessions that could be unambiguously mapped to a current Ensembl gene ID and HUGO Gene Nomenclature Committee (HGNC) gene symbol [33] via TogoID (accessed on August 9, 2025) [34] were retained to ensure gene-level consistency for downstream interpretation. To identify hit pairs with contrasting levels of sequence and structural similarity, the retained rice–human hit pairs were classified using two metrics: (i) global pairwise sequence similarity (%) computed by EMBOSS Needle (global alignment, default parameters) and (ii) the average lDDT score for the structural similarity metrics. Percentile thresholds (Q2 = median; Q3 = 75^th^ percentile) were computed across all retained pairs after filtering:

Low-sequence similarity/High-structural similarity (LS–HS): sequence similarity ≤ Q3 and average lDDT ≥ Q2.

High-sequence similarity/High-structural similarity (HS–HS): sequence similarity > Q3 and average lDDT ≥ Q2.

The sequence similarity cutoff at the 75^th^ percentile (Q3) was selected to retain a sufficient number of LS–HS pairs for downstream analyses while separating high-similarity matches into HS–HS.

Finally, “unique LS–HS” pairs were defined as LS–HS pairs with a rice gene lacking an HS–HS pair; LS–HS pairs from rice genes that had at least one HS–HS pair were excluded from the “unique” set.

### 2.6. Validation of Hit Pairs Using Other Public Database Resources

To compare the selected structural hit pairs with existing knowledge, two types of information from public databases were integrated.

1. Domain information: For each rice and human protein, InterPro domain IDs were retrieved via UniProt (accessed August 9, 2025) [28] [35], to confirm that they shared a common protein domain.
2. Orthologous relationships: Using rice gene IDs as a query, the Ensembl REST API (Comparative Genomics; Pan-taxonomic compara) [36] (accessed May 24, 2025) was used to identify orthologous relationships between each rice gene and its human counterpart (API example: https://rest.ensembl.org/homology/id/oryza_sa-tiva/Os01g0180800?compara=pan_homology&content-type=application/json;target_taxon=9606).

### 2.7. Workflow Implementation

To generalize and expand the analysis process implemented in sections 2.5, the “plant2human workflow” was developed, as described in the Common Workflow Language (CWL) [37], [38]. This series of processes ultimately produced a Jupyter notebook as an analysis report, allowing users to refine the filtering criteria and adopt additional actions as necessary. This workflow is available for free download from the WorkflowHub website under an MIT License [39].

## 3. Results

### 3.1. Functional Annotation Workflow

Figure 1 outlines the three-step functional annotation workflow of this study. First, a meta-analysis of rice RNA-Seq data under heat stress identified the upregulated and downregulated genes. Second, the predicted protein structures and sequences from these genes were aligned between rice and humans to identify pairs with low sequence similarity and high structural similarity (unique LS–HS condition) between rice and humans. Third, the selected hit pairs were further investigated using other public databases.

**Figure 1.**
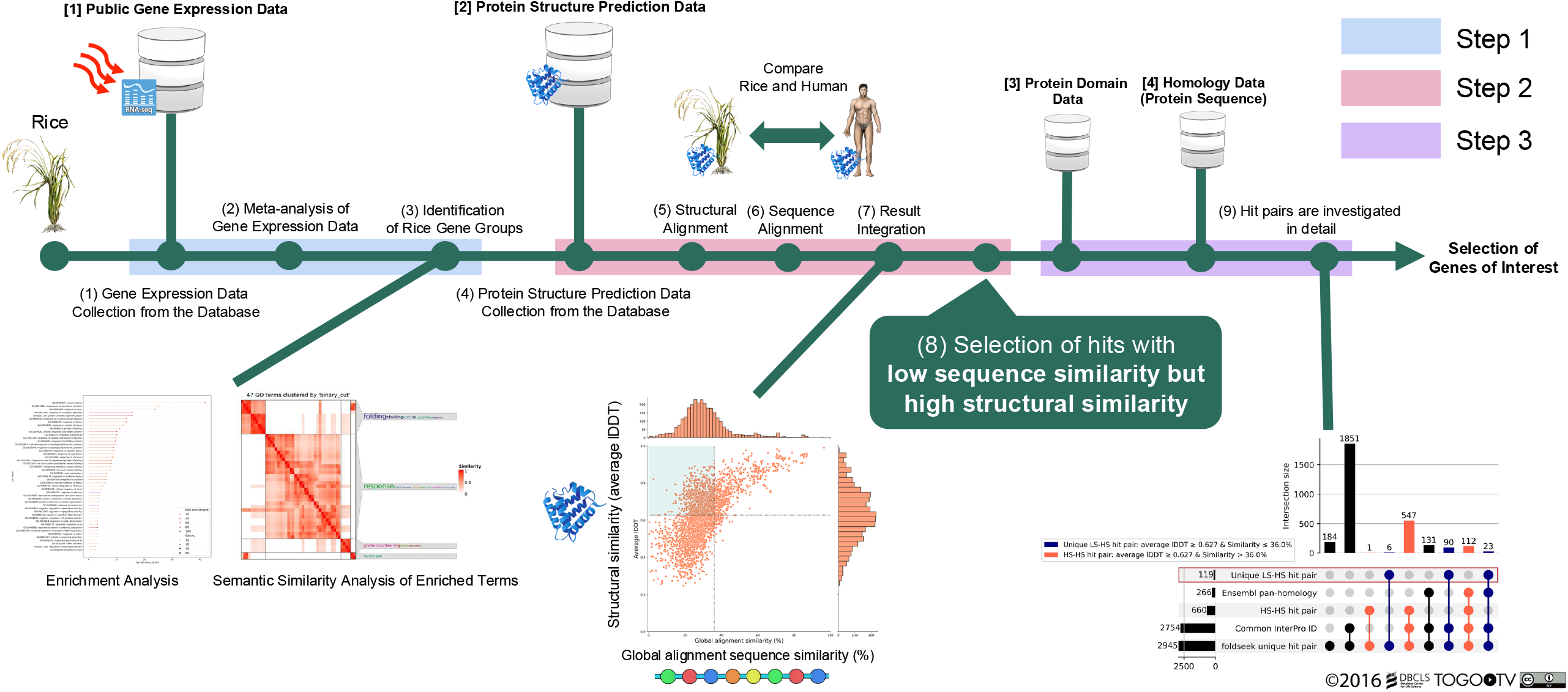
Summary of the functional annotation workflow in rice heat stress-responsive genes.

### 3.2. Gene Group Evaluation

We manually curated 360 pairs (heat stress vs. control conditions) of rice heat stress-related RNA-Seq data from 13 projects in the NCBI GEO (Fig. 1). The collected data comprised a wide range of experimental conditions, including heat shock and prolonged exposure to heat stress (e.g., high night temperature stress). Background information was available as a manually collected metadata CSV file (Supplementary Table S1) [40]. Background information with complex relationships was visualized using a Sankey diagram (Supplementary Fig. S1 [40]).

Subsequently, expression quantification, HN-ratio calculation (expression ratio), and evaluation index (HN-score) calculation were performed on the collected rice RNA-Seq data (Supplementary Tables S2–4 [40]). Based on the HN-score, rice genes within the top 1% (42 ≤ HN-score ≤ 255; 367 genes) and bottom 1% (−204 ≤ HN-score ≤ −40; 370 genes) were selected as the upregulated and downregulated gene groups, respectively (Fig. 2A, Supplementary Table S5) [40].

**Figure 2.**
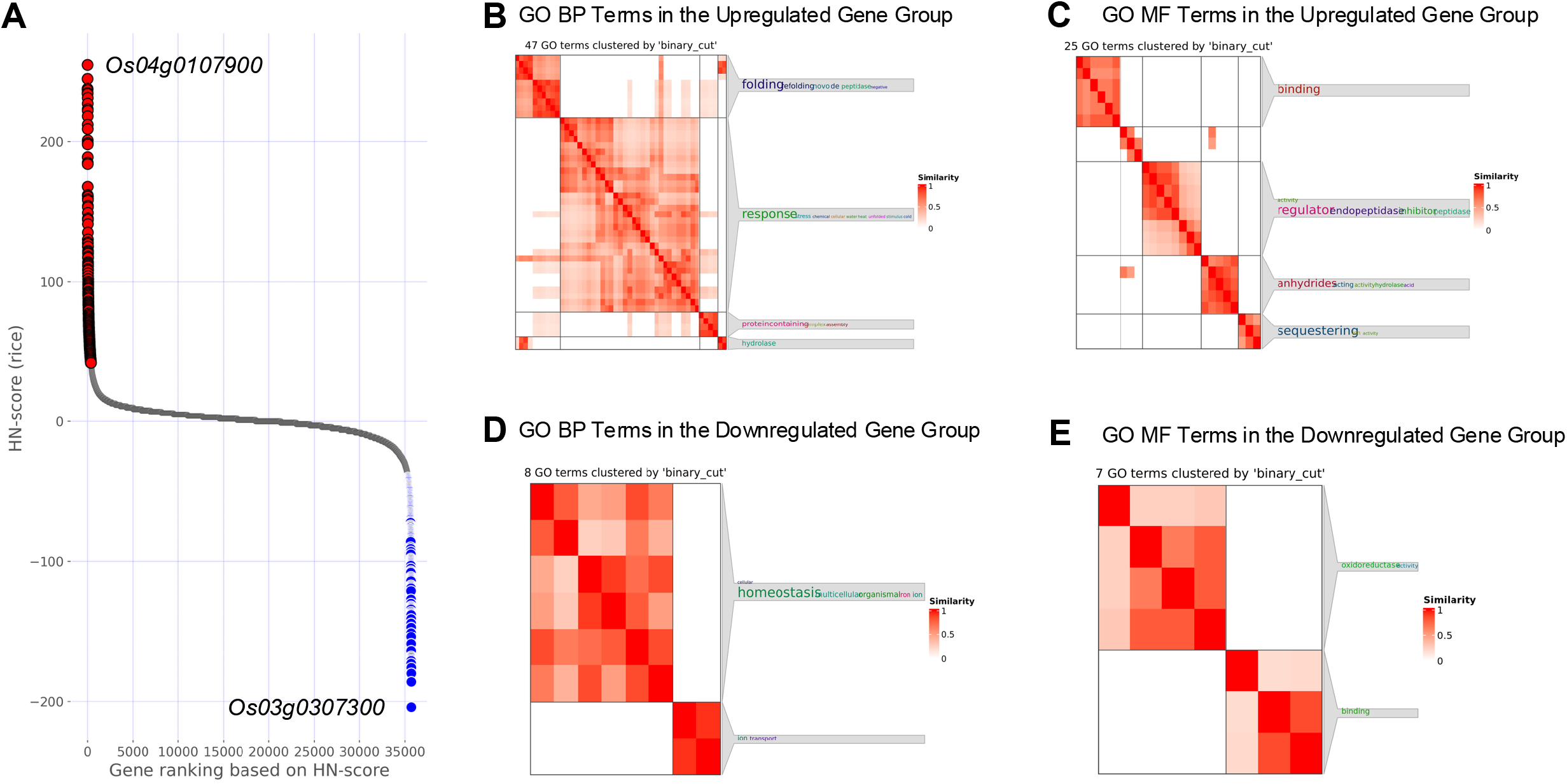
Characteristics of the gene groups selected by the HN-score. (A) Scatter plot of HN-scores for all rice genes; red: upregulated (*n* = 367), blue: downregulated (*n* = 370). Genes shown at the top left and bottom right indicate genes with the highest (255) or lowest (−204) HN-scores, respectively. (B, C) Clustering results of terms enriched in the upregulated group: (B) BP; (C) MF. Clustering results of terms enriched in the downregulated group: (D) BP, (E) MF. CC clusters are shown in Supplementary Fig. S3 [40]. Abbreviations: GO, Gene Ontology; BP, Biological Process; MF, Molecular Function

GO enrichment analysis (Supplementary Fig. S2 and Supplementary Table S6 [40]) were performed to characterize both groups, and the enriched terms were clustered by semantic similarity using simona [24] (Fig. 2B–E; Supplementary Fig. S3) [40]. The upregulated gene group included GO:0009408, “response to heat” (Fig. 2B), and GO:0051082, “unfolded protein binding” clusters (Fig. 2C), aligning with our Step 1 curation and HN-score ranking by effectively capturing heat-stress-responsive genes. In the downregulated gene group, a cluster related to homeostasis (e.g., GO:0098771, inorganic ion homeostasis; Fig. 2D) was detected, as well as a separate cluster including GO:0016491 oxidoreductase activity (Fig. 2E). Clusters associated with metal ion-related metabolism—especially iron—were observed: the upregulated group included GO:0140315 iron ion sequestering activity (such as *Os11g0106700, Os12g0106000*, and *Os09g0396900*), whereas the downregulated group included GO:0006826 iron ion transport (such as *Os02g0649900*) (Fig. 2C, D). Furthermore, the term enriched in the upregulated gene group included GO:0046688 response to copper ion (*Os03g0266900, Os03g0267000*, and *Os03g0267200*) (Supplementary Fig. S2 and Supplementary Table S6 [40]).

### 3.3. Structurome Analysis

Using the gene lists described in Section 3.2, the predicted rice protein structures were compared with those of humans, focusing on LS–HS candidates by combining structural and sequence information (Step 2 in Fig. 1). Pair-wise sequence alignment results, together with structural alignment data, were used to identify protein hit pairs across distantly related species.Predicted structures were obtained from the AFDB, and structural searches were conducted using Foldseek (3Di+AA and Foldseek-TM method) against the complete AFDB database [6], [29]; rice–human hits were retained (Supplementary Table S7 [40]). The 3Di+AA results served as the primary analysis; the Foldseek-TM findings are summarized in Supplementary Fig. S4 and Supplementary Table S8 [40]. After pairwise sequence alignment and the application of predefined filters, 2,945 hit pairs were identified in the upregulated gene group (145/367 genes) and 3,708 hit pairs in the downregulated gene group (147/370 genes). The distributions of the average lDDT and global sequence similarity (%) are shown in Fig. 3A and B, with additional metrics and indicators in Supplementary Fig. S5, and all hit pairs in Supplementary Table S9 [40].

**Figure 3.**
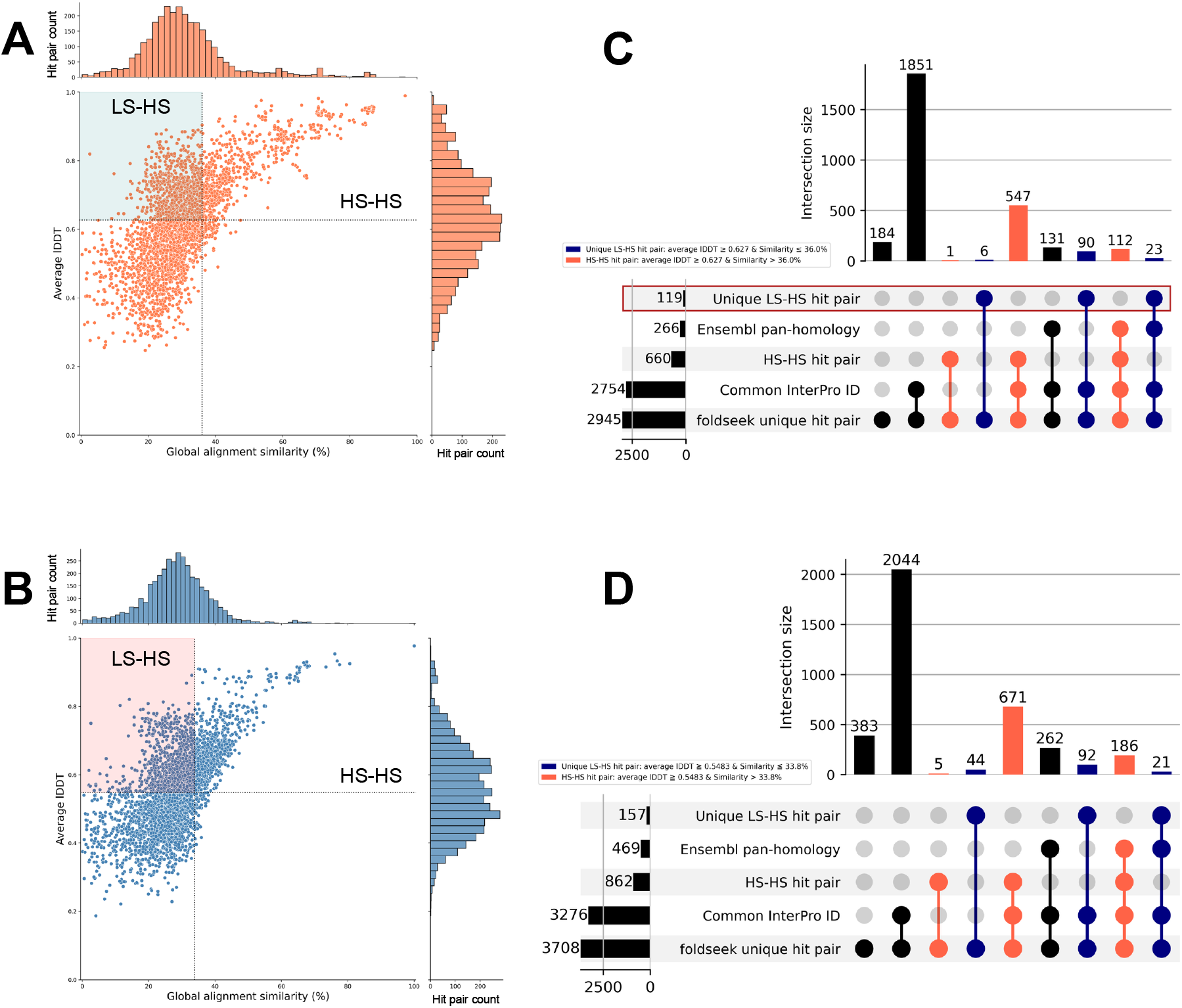
Landscape of rice–human structural/sequence relationships and characterization of hit pairs using public database information. (A, B) Scatter plots of average lDDT (y-axis) versus global pairwise sequence similarity (%) computed using EMBOSS Needle (x-axis) for the retained rice–human pairs. Vertical and horizontal lines denote Q3 (sequence similarity) and Q2 (average lDDT), respectively: (A) derived from the upregulated gene list. Vertical line Q3 = 36.0%; horizontal line Q2 = 0.627; (B) derived from the downregulated gene list. Vertical line Q3 = 33.8%; horizontal line Q2 = 0.5483. (C, D) UpSet plots showing intersections among five attributes based on hit pair count. Rows (bottom to top): Foldseek all unique hit pairs, Common InterPro ID (Share at least one InterPro ID), HS–HS hit pair (sequence similarity > Q3, average lDDT ≥ Q2), Ensembl pan-homology (ortholog relationships based on protein sequence), and Unique LS–HS hit pair (sequence similarity ≤ Q3, average lDDT ≥ Q2). Left horizontal bars indicate the set size of each row. Columns indicate set intersections; closed circles denote membership, and vertical connectors denote logic. The bar height at the top shows the intersection size (number of hit pairs); blue: intersection consists exclusively of “Unique LS–HS” hit pairs, red: intersection consists exclusively of “HS–HS” hit pairs, black: other combinations. (C) upregulated; (D) downregulated.

Within the LS–HS regions (Fig. 3A, B), 119 hit pairs were identified (38/367 genes) in the upregulated gene group and 157 hit pairs (34/370 genes) in the downregulated gene group after excluding rice genes that also had HS–HS matches (“unique LS–HS”). Subsequently, “unique LS–HS” and HS–HS pairs were compared using InterPro domain information and ortholog information between rice and humans retrieved from Ensembl (Fig. 3C, D; Step 3 in Fig. 1) [19], [35]. By combining these two types of information, we identified hit pairs in which existing data (domain and ortholog information or domain information alone) supported the correspondence of rice–human structural hit pairs or cases in which both types of information were missing (Fig. 3C, D). To demonstrate specific examples, six representative rice–human unique LS–HS hit pairs were evaluated (Table 1) using different combinations. These genes did not appear in the GO enrichment analysis (Supplementary Fig. S2, and Table S6 [40]). All hit pairs and details of unique LS–HS and HS–HS interactions are presented in Supplementary Table S9 [40].

**Table 1.**
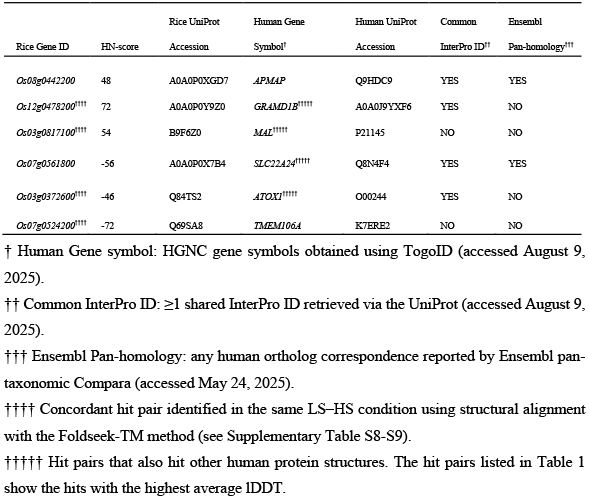
Representative unique LS–HS rice–human hit pairs.

## 4. Discussion

This study presents an analytical workflow integrating public transcriptome and structural data to investigate heat stress response genes in rice.

### 4.1. Heat Stress Response System and Metal Metabolism

Micronutrient metals such as iron and copper are essential for the survival of land plants, including rice. The molecular mechanisms responsible for regulating metal homeostasis and the associated genes are well characterized [41]. Through a meta-analysis of publicly available transcriptome data under heat stress conditions and accompanying structural analysis, genes related to metal homeostasis were identified.

Enrichment analysis with Gene Ontology (GO) and semantic similarity analysis revealed that genes involved in iron ion sequestration were upregulated, whereas those involved in iron ion transport were downregulated in rice (Supplementary Table S6 [40]; Fig. 2C, D). These changes may reflect characteristics observed in ferroptosis under heat stress. The upregulated gene set included *Os12g0106000* (gene symbol: *OsFER2*), which positively regulates ferroptotic cell death when rice is infected with avirulent *Magnaporthe oryzae* as a form of biotic stress [42]. Furthermore, in *A. thaliana*, regulated cell death under heat stress occurs in the form of ferroptosis-like cell death [43]. However, direct evidence explaining the coordinated regulation of these gene sets under heat stress remains limited. Recent research has explored the interaction between temperature stress and iron. For instance, studies on light-chilling stress in cucumbers (*Cucumis sativus L*.) demonstrate that high-iron nutritional conditions can exacerbate oxidative damage via light-dependent root-to-shoot iron translocation and lipid peroxidation, whereas low-iron conditions may mitigate these effects [44]. As such, insights into iron nutrition management have direct implications for agricultural applications [44]. Enrichment and structural analyses in humans identified genes potentially related to copper and iron metabolism. The upregulated gene group was enriched for the GO:0046688 response to copper ion (Supplementary Fig. S2 and Supplementary Table S6 [40]). Structural comparisons with humans highlighted gene pairs like *Os03g0372600–ATOX1* (Table 1, row 5), sharing the heavy metal-associated InterPro domain (IPR006121, accessed August 21, 2025). Although A rice gene belonging to the same domain (IPR006121), *Os02g0196600* (*OsHMA4*), is known to transport copper to root vacuoles [45], *Os03g0372600* remains understudied [46]. Insights from human copper counterparts—*ATOX1* (antioxidant 1 copper chaperone)—and the emerging concept of cuproptosis—copper-dependent cell death [47]—can inform hypothesis generation for heat-stress response systems in rice.

High temperatures affect rice throughout its life cycle and across various plant organs. Genes linked to heat tolerance have been associated with plant growth stages, organs, and thermophenotypes [13]. Further investigation into metal metabolism-related genes identified in this study, particularly their thermophenotypes, could aid in future agricultural applications.

### 4.2. Comparison with Distantly Related Species through Structural Similarity

As a proof-of-concept, we aimed to identify novel heat stress-responsive genes using public transcriptomes and structures. Our selection of humans as the target organism was motivated by the opportunity to utilize conserved protein structural features, rather than strict orthology, to map rice proteins into a comprehensively annotated human context [9]. For instance, hit pairs with high structural similarity but uneven annotations, such as *Os03g0372600*–*ATOX1* pair (Table 1, row 5), enable the transfer of human annotations to rice, generating testable hypotheses.

Additionally, our findings suggest that conserved protein structure can support the use of rice as a model organism for humans—a concept consistent with prior validations of *BRCA2* in *Arabidopsis* and confirmation of human disease gene orthologs in *Arabidopsis* and *P. trichocarpa* [10], [11]. With resources like the AFDB now offering large-scale protein structure datasets [6], leveraging conserved protein structures presents a scalable approach for nominating a broader range of species as hypothesis-generating model organisms relevant to human research. For example, metal metabolism highlighted in section 4.1 underscores the translational potential of plant findings for human biology. Such comparative analyses are facilitated by the “plant2human workflow,” implemented in CWL [37], [38], which allows systemic mapping between genes in various species and their human counterparts via the structurome. This workflow enhances biological discovery by enabling reciprocal functional annotation across species.

To investigate distant functional protein hit pairs that may not be detected by sequence alignment-based methods, low (global) sequence similarity combined with high structural similarity (LS–HS condition) was used in conjunction with three filtering conditions. This methodology draws upon the concept of the “twilight zone,” a region in sequence alignment where homology inference is difficult due to sequence identity limitations [48]. Recent approaches targeting proteins in the twilight zone use physicochemical property-based methods [49]. In this study, the approach was extended by introducing an exploratory “user-defined twilight zone” referred to as the LS–HS condition to facilitate detection of understudied genes. Sequence similarity served as a general indicator rather than an optimization target for structural hit pairs; thus, sequence-alignment parameters were maintained at default values (see Section 2.5.3). Although further refinement could enhance the method, it is likely to contribute new perspectives in evaluating individual genes, including those excluded from enrichment analyses.

The obtained hit pairs were cross-referenced with the InterPro domain data and Ensembl ortholog information to identify currently available data from the protein sequences. This integration classified hit pairs, that will help prioritize rice genes of interest. For example, hit pairs sharing common domains and orthologous relationships may be supported by structural and sequence evidence, potentially enabling the annotation of human proteins to serve as a starting point for the functional analysis of rice genes. Conversely, hit pairs lacking domain information and ortholog correspondence, that is, hit pairs not supported by sequence information, are highly intriguing as protein hit pairs supported solely by structural details but require careful interpretation. Furthermore, continuous updating of the sequence information recorded in UniProt could lead to hit pairs with discrepancies between structural and domain information. Therefore, there are hit pairs that require actions, such as reacquiring domain information and using AlphaSync, which synchronizes protein structure data [50].

### 4.3. Limitation

This study has several methodological limitations that could lead to misannotation. First, we did not impose stringent quality-based filtering on AlphaFold predictions (e.g., predicted Local Distance Difference Test (pLDDT) thresholds [5]), prioritizing coverage to consider more genes. This choice increases sensitivity but may also lead to inflated false-positive structural matches, especially in regions of low confidence. Second, our comparison focused on more global structural similarities and did not evaluate detailed structural verification, such as that performed for the rice HSP90 protein structure [15]. Therefore, interpreting gene functions still requires careful consideration, necessitating experimental verification similar to that performed in previous studies that identified unknome [12]. In summary, for understudied genes identified in this research, additional verification using genome editing technologies is necessary to prevent misannotation.

## 5. Conclusion

This study focuses on poorly understood heat stress-responsive genes in rice by integrating resources from public life science databases, particularly public transcriptome and structural data. This approach identifies understudied genes and suggests the potential for functional annotation using information from other species, such as humans, through structurome analysis. Furthermore, the newly developed plant2human workflow is effective for heat stress and exploring gene groups with diverse biological backgrounds.

## Acknowledgements

Computations were performed on computers at the Hiroshima University Genome Editing Innovation Center.

## Author contributions

Conceptualization: S. Y. and H. B.; methodology: S. Y. and H. B.; software: S. Y.; validation: S. Y. and H. B.; formal analysis: S. Y.; investigation: S. Y.; resources: H. B.; data curation: S. Y.; writing—original draft preparation: S. Y.; writing—review and editing: S. Y. and H. B.; visualization: S. Y.; supervision: H. B.; project administration: H. B.; funding acquisition: H. B. All authors have read and agreed to the published version of the manuscript.

## Supplementary data

All supplementary information is available on the figshare website (https://doi.org/10.6084/m9.figshare.c.7835495.v3) [40].

## Data availability

All codes in this study are publicly available at https://github.com/yonesora56/HS_rice_analysis and are licensed under the MIT license.

This analysis workflow, executed in sections 2-5, is available as a “plant2human workflow” in WorkflowHub, licensed under the MIT license [37], [39].

